# Crucial roles of mesenchymal *Gata2* in murine epididymal development

**DOI:** 10.1101/2025.03.26.645498

**Authors:** Allyssa Fogarty, Shuai Jia, Jillian Wilbourne, Claire DuPuis, Fei Zhao

## Abstract

Androgens drive the morphogenesis and differentiation of the Wolffian duct (WD) into the epididymis, an essential organ for male reproduction, by binding to the androgen receptor (AR). However, it remains unclear whether other transcriptional programs operate beyond the central androgen/AR signaling in promoting WD development. We discovered that mesenchyme-specific deletion of the transcription factor *Gata2* resulted in defective epididymal coiling in the corpus and caudal regions. The defective coiling in the absence of mesenchymal *Gata2* did not result from androgen signaling deficiency, as there were no abnormalities in testicular morphology, androgen production, or AR/*Ar* expression, and dihydrotestosterone supplementation did not restore epididymal coiling in cultured WDs. Instead, *Gata2* deletion reduced the expression of the mesenchyme-derived factor *Inhba* and epithelial proliferation, both of which play critical roles in epididymal coiling. The epididymal defect persisted into adulthood, with the uncoiled corpus and caudal epididymis exhibiting abnormal epithelial morphology and lumen environments, resulting in an unfavorable environment for sperm storage. Our results demonstrate the androgen-independent role of mesenchymal GATA2 in promoting epididymal development through *Inhba* induction and highlight the importance of proper fetal development in male reproduction.

**Significance Statement:** Testicular androgens drive the maintenance and differentiation of the Wolffian duct into a coiled and functional epididymis during male sexual differentiation. Our study, however, reveals that the Wolffian duct failed to develop into a coiled and functional epididymis in the absence of mesenchymal *Gata2.* This defect is not due to androgen signaling deficiency but rather to reduced expression of the mesenchyme-derived factor *Inhba* and decreased epithelial proliferation. Our results demonstrate the crucial role of GATA2-mediated transcriptional programs beyond the central androgen/androgen receptor signaling in Wolffian duct development.

## Introduction

Alfred Jost’s landmark experiments in the 1940s and 1950s laid the foundation for our current paradigm of male sexual differentiation (1, 2). His work demonstrated that testicular androgens are crucial for male reproductive tract maintenance and differentiation. He showed that castrated male embryos lost the Wolffian duct (WD), the precursor to the male reproductive tract organs; however, androgen replacement could restore WD maintenance and promote its differentiation. The identification and cloning of the androgen receptor (AR) advanced our understanding of how androgen signaling is mediated (3). Subsequent studies on loss-of-function mutations of AR*/Ar* genes in both mice and humans have underscored the crucial role of androgen-AR signaling in WD development (4). Disruption in androgen signaling by androgen receptor antagonists during the fetal masculinization programming window can lead to various adverse effects on WD development, including incomplete and underdeveloped epididymides (5, 6). Thus, accumulating evidence has solidified the paradigm that androgen-AR signaling is the primary driver for WD development.

Development of the WD is primarily driven by androgen-AR signaling within the mesenchyme, which is the undifferentiated connective tissue that surrounds the epithelium. Classical tissue recombinant studies have demonstrated the instructive role of the mesenchyme in epithelial morphogenesis and differentiation. Under the influence of androgen action, the upper, middle and lower WD differentiates into the epididymis, vas deferens and seminal vesicle, respectively (7). In the heterotypic tissue recombinant consisting of the upper and middle WD epithelium with lower WD mesenchyme, the epithelium underwent seminal vesicle morphogenesis and synthesized seminal vesicle-specific secretory proteins (8). Tissue-specific *Ar* ablation studies also demonstrate the crucial role of mesenchymal androgen/AR actions in WD development. Deletion of epithelial *Ar* did not impair WD maintenance or morphogenesis (9). However, mesenchyme-specific *Ar* deletion resulted in caudal WD degeneration and cystic formation in the cranial WD region (10). Collectively, these studies demonstrate that the mesenchyme governs the fate and differentiation of WD development in the male embryo.

Our current project aimed to uncover the critical role of mesenchymal GATA2 in promoting WD development. GATA2, a member of the GATA family of transcription factors, plays crucial roles in cell differentiation and organ development (11–13). Our published RNA-seq analysis of WD mesenchyme during fetal sexual differentiation identified *Gata2* as the highest expressed GATA transcription factor in the WD mesenchyme (14). Global knockout of *Gata2* leads to embryonic lethality at embryonic day (E)10.5 due to defective hematopoiesis, thereby limiting our investigation into GATA2’s role in the male reproductive tract development (15). To address this limitation, we used our established mesenchyme-specific *Osr2-Cre* (10) to study the mesenchyme-specific role of *Gata2* in WD development.

## Results

### Mesenchymal *Gata2* is critical for establishing epididymal coiling and identity

We first performed GATA2 immunofluorescent staining on cross-sections of the mesonephros in the cranial and caudal regions to determine its expression during sexual differentiation of the male reproductive tracts. At the time of WD stabilization (E14.5), GATA2 was localized to the nuclei of the mesenchymal cells surrounding both WD and Müllerian duct (MD) epithelium in both cranial and caudal regions of mesonephroi **(Fig. 1A)**. At E16.5, when the MD degenerates and the stabilized WD starts to undergo coiling, GATA2 was continuously expressed in the nuclei of mesenchymal cells in both regions (**Fig. 1A**).

**Figure 1.**
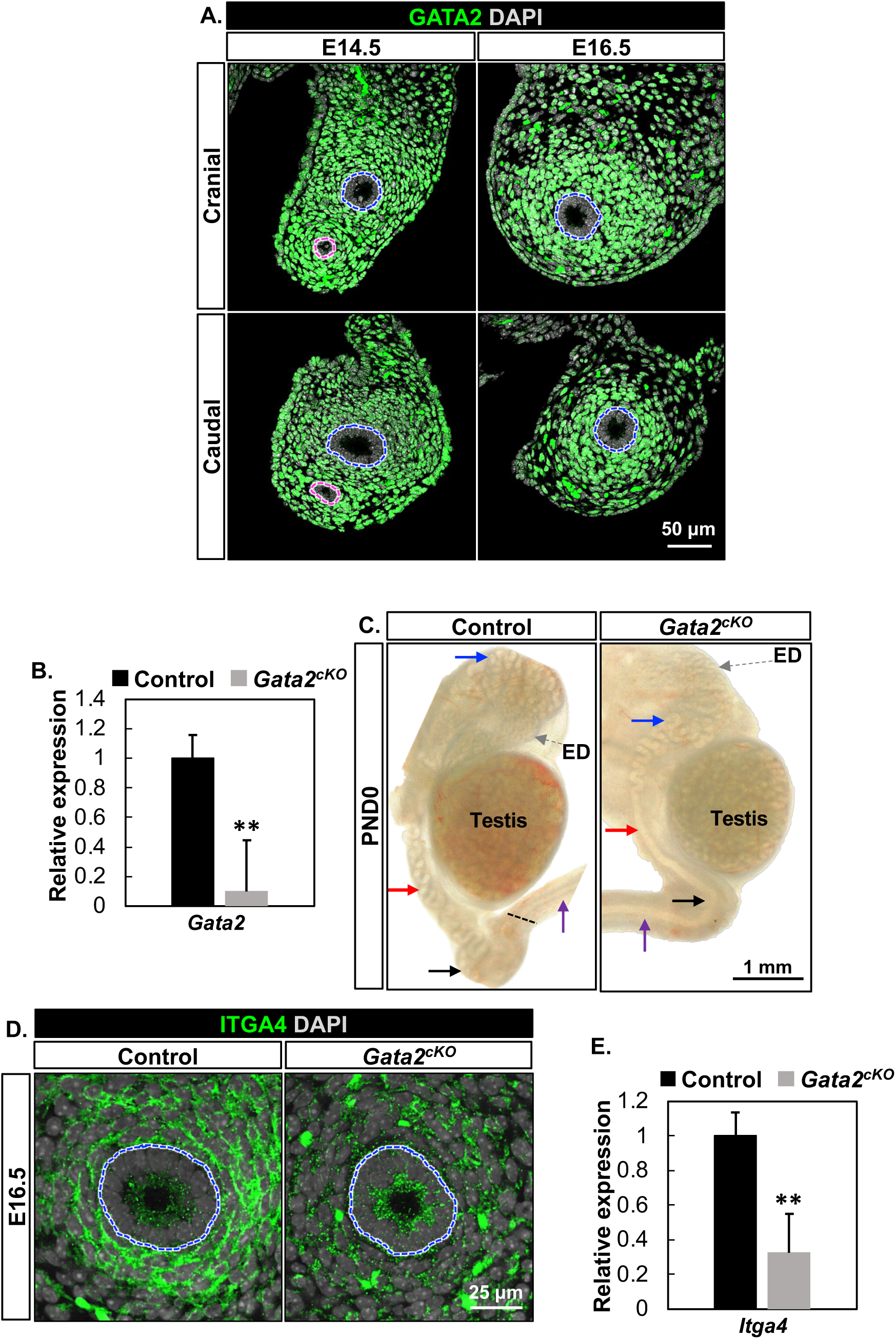
Mesenchymal *Gata2* is critical for establishing epididymal coiling and identity. **(A)** Immunofluorescent staining of GATA2 on cross-sections of the control mesonephroi in the cranial and caudal regions at E14.5 and E16.5 with n=4 for each timepoint. Blue and pink dashed lines: WD and Müllerian duct (MD), respectively. **(B)** Relative expression of *Gata2* mRNA in control and *Gata2^cKO^* mesonephroi at E16.5 with n=5 in each group. **(C)** Bright-field images of the control and *Gata2^cKO^* epididymides attached with testes at PND0 with n=3 in each group. Blue, red, and black arrows indicate cranial, corpus, and caudal regions, respectively. The grey and purple arrows: efferent ducts (ED) and the vas deferens, respectively. The dashed black line represents the border between the epididymis and vas deferens. **(D)** Immunofluorescent staining of ITGA4 on cross-sections of the control and *Gata2^cKO^* mesonephroi at E16.5 with n=4 in each group. Blue dashed lines: WD. **(E)** Relative expression of *Itga4* mRNA between the control and *Gata2^cKO^* mesonephroi at E16.5 with n=5 in each group. Results were shown as mean ± SEM and analyzed by 2-tailed *t* test. ** *P* < 0.01.

To determine the specific function of mesenchymal GATA2, we established a mesenchyme-specific conditional *Gata2* knockout mouse model (*Osr2-Cre:Gata2-flox)*. The successful deletion of *Gata2* in the *Gatat2^cKO^*mesonephori was confirmed by RT-qPCR **(Fig. 1B)**. At birth, control WDs became highly coiled throughout the cranial, corpus, and caudal regions of the epididymis (**Fig. 1C**). In the caudal region, we observed a discernible border between the coiled epididymis and the straight vas deferens due to their distinct morphologies **(Fig. 1C)**. However, the WDs in *Gata2^cKO^*newborns failed to elongate and coil in the corpus and caudal regions, forming a straight duct similar to the vas deferens. As a result, the border between the epididymis and vas deferens became indistinguishable **(Fig. 1C)**.

Since the corpus and caudal regions of the epididymis lost their characteristic coiling and resembled a straight vas deferens, we were curious whether the morphological change was accompanied by the loss of epididymal identity. We examined the expression of ITGA4/*Itga4*, a gene enriched in the epididymal region of fetal WDs (16). In the control group, ITGA4 was robustly expressed in both the membrane and cytoplasm of control WD mesenchymal cells at E16.5. However, its expression was significantly reduced in the *Gata2^cKO^* mice **(Fig. 1D & 1E)**, suggesting a loss of epididymal mesenchymal identity in *Gata2^cKO^* WDs. Taken together, our results demonstrate that GATA2 in the mesenchyme plays a significant role in establishing epididymal coiling and identity.

### Androgen signaling was unaffected in *Gata2^cKO^* male embryos

The defective WD coiling pointed to a possible deficiency in testicular androgen signaling, which is the predominant driver for WD development (17, 18). To test this possibility, we first examined testicular histology and expression of steroidogenic enzymes. We found comparable testicular cord structures between control and *Gata2^cKO^* males **(Fig. 2A)**. The expression of *Cyp17α1* and *Hsd3b1* (19), two rate-limiting steroidogenic enzymes required for androgen production, were comparable between control and *Gata2^cKO^* testes at E16.5 **(Fig. 2B).** These results indicated normal testicular development and androgen synthesis in *Gata2^cKO^* males.

**Figure 2.**
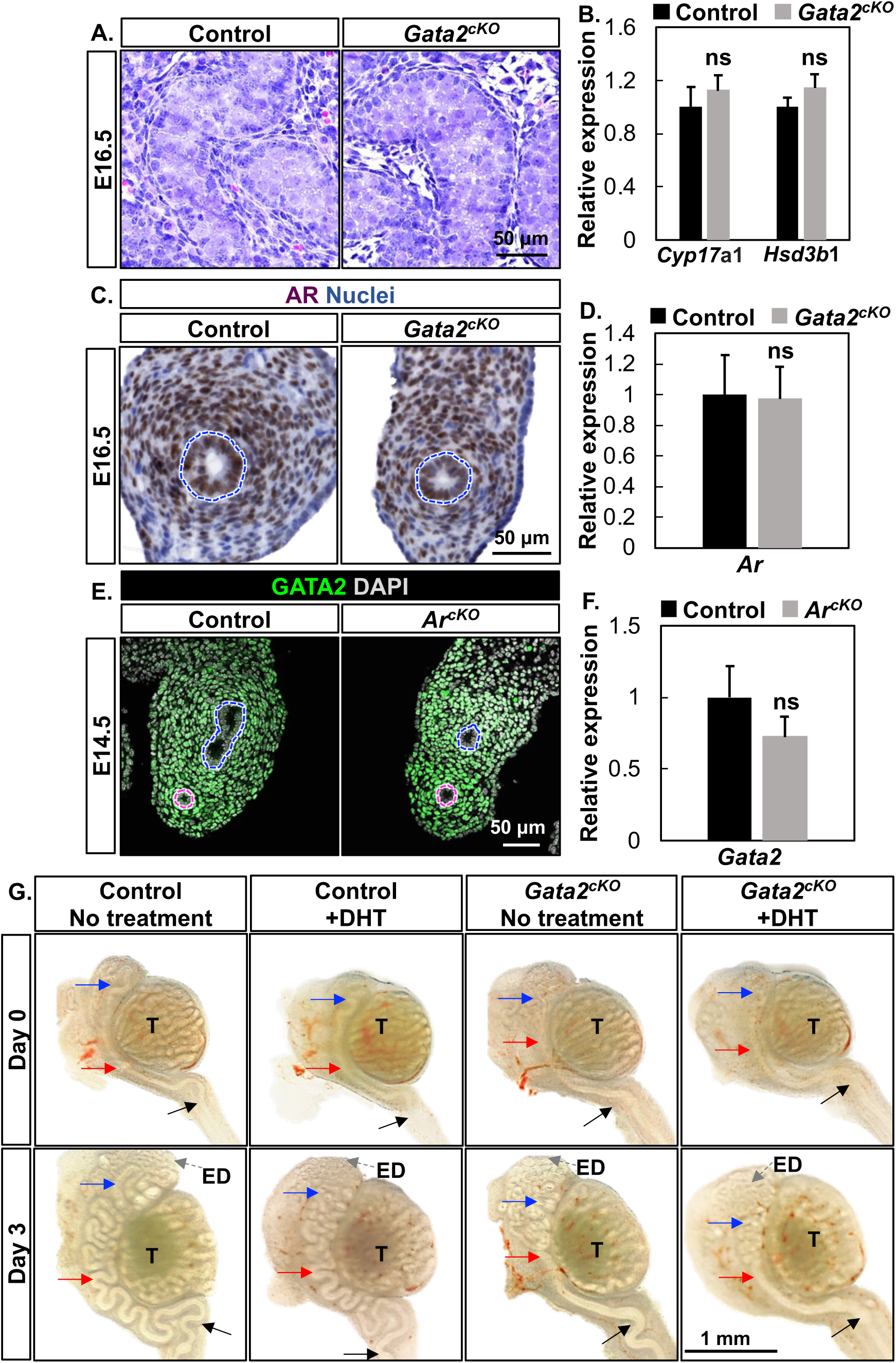
Androgen signaling was unaffected in *Gata2^cKO^* male embryos. **(A)** Hematoxylin and Eosin staining of the control and *Gata2^cKO^* testes at E16.5 with n=3 in each group. **(B)** Relative expression of the two steroidogenic enzymes *Cyp17a1* and *Hsd3b1* by RT-qPCR in the control and *Gata2^cKO^* testes at E16.5 with n=3 in each group. **(C)** Immunohistochemistry staining of AR on cross-sections of the control and *Gata2^cKO^* mesonephroi at E16.5 with n=4 in each group. Blue dashed lines: WD. **(D)** Relative expression of *Ar* mRNA between control and *Gata2^cKO^* mesonephroi at E16.5 with n=5 in each group. **(E)** GATA2 immunofluorescent staining on cross-sections of the control and *Ar^cKO^*mesonephroi at E14.5 with n=4 in each group. Blue and pink dashed lines: WD and MD, respectively. **(F)** Relative expression of *Gata2* mRNA in control and *Ar^cKO^* mesonephroi at E14.5 with n=5 in each group. Results in (B, D, F) were shown as mean ± SEM and analyzed by 2-tailed *t* test; ns = not significant. **(G)** Organ culture of E16.5 control and *Gata2^cKO^* mesonephroi with testes for three days with no treatment or DHT with n=3 for each group. “T” marks the testis; the blue, red, and black arrows indicate cranial, corpus, and caudal regions, respectively; the grey dashed arrows: ED.

The androgen action is mediated by the AR in male reproductive tract organs.

Thus, we examined AR/*Ar* expression in the mesonephroi using immunohistochemistry and RT-qPCR. AR was detected in the nuclei of the epithelium and its surrounding mesenchyme in both control and *Gata2^cKO^* mesonephroi **(Fig. 2C)**. *Ar* expression at the mRNA level was also comparable between control and *Gata2^cKO^*mesonephroi **(Fig. 2D)**. These findings indicated that AR/*Ar* expression was unaffected in the absence of mesenchymal *Gata2*.

Another possible involvement of androgen signaling is that *Gata2* might be a downstream gene of the mesenchymal AR. To determine this possibility, we examined GATA2/*Gata2* expression in the mesonephroi of our previously established mesenchyme-specific *Ar* knockout mouse model (*Ar^cKO^*) (10). GATA2 immunofluorescent staining showed nuclei staining in the mesenchymal cells surrounding WDs and MDs between control and *Ar^cKO^* mesonephroi at E14.5 **(Fig. 2E)**. The expression of *Gata2* at the mRNA level was comparable between control and *Ar^cKO^* mesonephroi by RT-qPCR **(Fig. 2F)**. These results demonstrated that *Gata2* does not act as a downstream factor of mesenchymal androgen/AR signaling.

To conclusively rule out that androgen signaling deficiency was the cause of the *Gata2^cKO^* phenotype, we supplemented cultured WDs and testes with the potent and non-aromatizable androgen, dihydrotestosterone (DHT), at the established concentration of 10nM (20). At the beginning of culture (E16.5), WDs in both control and *Gata2^cKO^* groups were straight with the initial coiling in the caput region **(Fig. 2G)**. After three-day culture, WDs from the control embryos underwent elongation and extensive coiling at the caput, corpus and caudal regions in both the no treatment and DHT treatment groups. By contrast, *Gata2^cKO^* WDs failed to become coiled in corpus and caudal regions, recapitulating our observed phenotype in vivo **(Fig. 2G)**. More importantly, *Gata2^cKO^*WDs in the DHT treatment group remained uncoiled in the corpus and caudal regions, demonstrating that androgen supplementation is not able to rescue the *Gata2^cKO^* phenotype. Collectively, our results demonstrated that the defective WD coiling in the *Gata2^cKO^* embryo is not due to deficient androgen signaling.

### The expression of the key mesenchyme-derived factor *Inhba* was reduced in *Gata2^cKO^*

*Inhba* (Inhibin βA) is the only known mesenchyme-secreted factor that governs epididymal coiling by promoting epithelial proliferation (21). We, therefore, assessed its expression using RNA-scope on cross-sections of control and *Gata2^cKO^* E16.5 mesonephroi **(Fig. 3A)**. *Inhba* expression was primarily localized in the mesenchyme surrounding WD epithelium in both control and *Gata2^cKO^* mesonephroi. However, *Inhba* expression was reduced in the mesenchyme of *Gata2^cKO^* mice **(Fig. 3A)**. The percentage of the mesenchymal cells expressing *Inhba* **(Fig.3B)** and the average expression score in those *Inhba+* cells **(Fig. 3C)** were both significantly reduced. A major mechanism by which mesenchymal *Inhba* regulates WD coiling is promoting epithelial cell proliferation (21). We therefore investigated whether epithelial cell proliferation was reduced in *Gata2^cKO^* mice by running immunofluorescent staining of cell proliferation maker KI67 **(Fig. 3D)**. We found the percentage of KI67+ epithelial cells was significantly reduced in the absence of mesenchymal *Gata2* **(Fig. 3E)**. These results demonstrated that *Inhba* is a potential downstream effector of mesenchymal GATA2 in regulating epididymal coiling.

**Figure 3.**
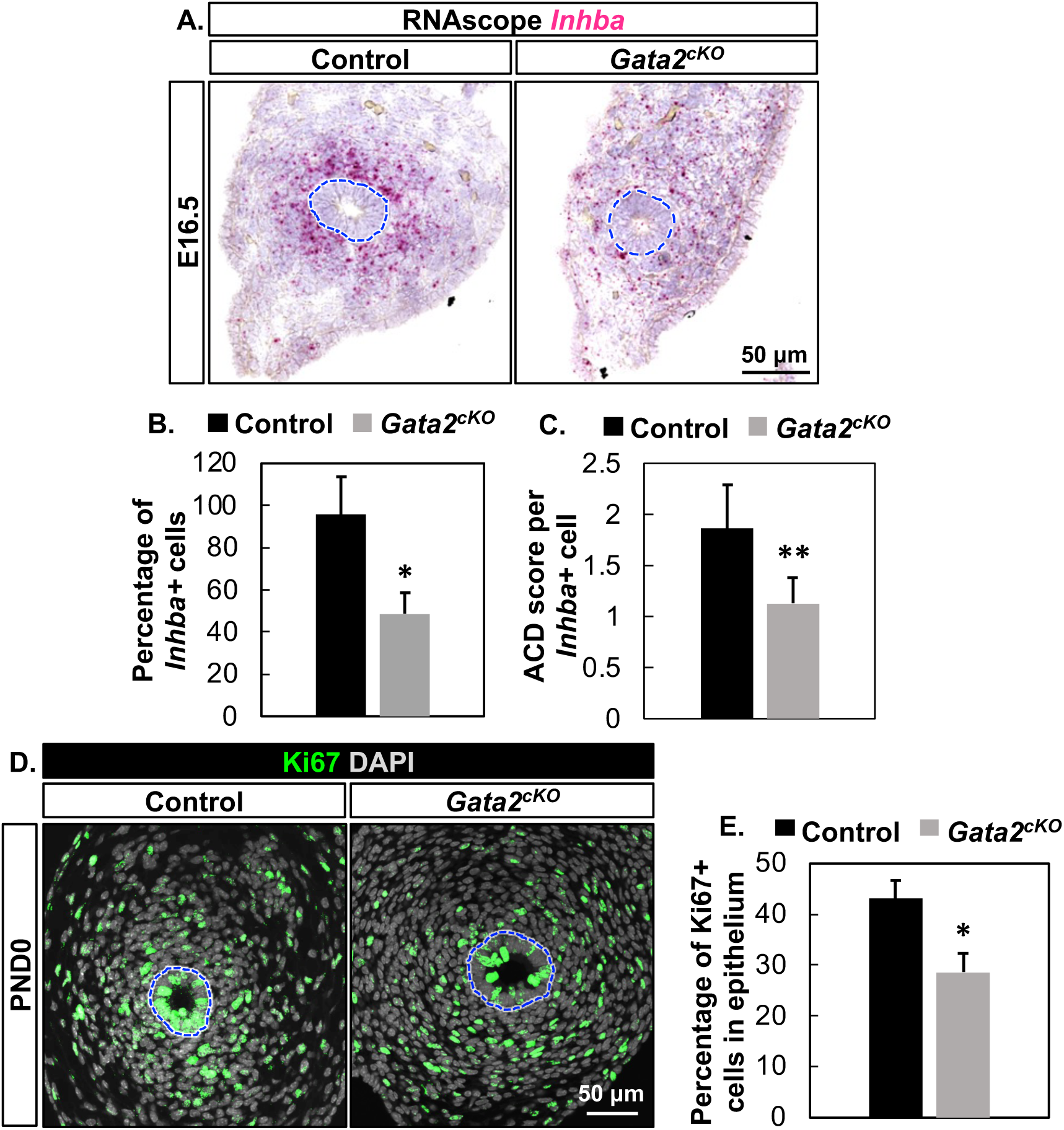
The expression of the mesenchyme-derived factor *Inhba* and WD epithelial proliferation were reduced in *Gata2^cKO^* embryos. **(A)** Representative images of *Inhba* RNA-scope staining on cross-sections of the control and *Gata2^cKO^* mesonephroi at E16.5 with n=4 embryos in each group. Blue dashed lines: WDs. **(B)** The average percentage of *Inhba+* cells in the mesenchyme with n=4 embryos in each group. **(C)** Average ACD score per *Inhba+* cell with n=4 embryos in each group. The average of results from four to eight sections per embryo was used for statistical analysis. **(D)** Immunofluorescent staining of KI67 on cross-sections of the control and *Gata2^cKO^* epididymides at PND0 with n=6 embryos in each group. Blue dashed lines: WDs. **(E)** The percentage of KI67+ epithelial cells in the control and *Gata2^cKO^* epididymides at PND0 with n=6 embryos in each group. The average of results from five to eight sections per embryo was used for the statistical analysis. Results were shown as mean ± SEM and analyzed by 2-tailed *t* test; ns=not significant; * *P* < 0.05; ** *P* < 0.01.

The major product of *Inhba* in the mesonephros has been suggested to be Activin A (homodimer of Inhibin βA), which activates type I and II receptors (22) and ultimately leads to activation of SMAD signaling in WD epithelium (21). We then were curious whether the SMAD-mediated signaling in WD epithelium was indispensable for WD morphogenesis. To address this question, we deleted the common mediator of SMAD signaling, *Smad4,* in the WD epithelium using *Hoxb7-Cre* (23). SMAD4 was expressed in the nucleus and cytoplasm in the entire epithelium of control groups. In *Smad4^cKO^* mesonephroi, many of the epithelial cells lost SMAD4 expression **(Fig.S1A)**. In both control and *Smad4^cKO^* male embryos, their epididymides were normally coiled **(Fig. S1B)**. These findings indicated that SMAD4-mediated signaling is dispensable for WD coiling, and that mesenchymal INHBA may act through SMAD4-independent pathway(s) in promoting WD coiling.

### Adverse consequences on differentiation and function of the adult epididymis

After birth, the epididymis continues to develop before becoming mature and functional in supporting post-testicular sperm transport, maturation and storage during puberty (7). To investigate the impact of the deletion of mesenchymal *Gata2* on the postnatal epididymis, we examined testes and epididymides from control and *Gata2^cKO^* males at the age of sexual maturity (8-weeks) **(Fig. S2A)**. The size and histology of testes between control and *Gata2^cKO^* groups were comparable with spermatogenesis in the lumen of normally looking seminiferous tubules **(Fig. S2B)**. However, *Gata2^cKO^* adult epididymides were smaller in size than those of the control **(Fig. 4A)**.

**Figure 4.**
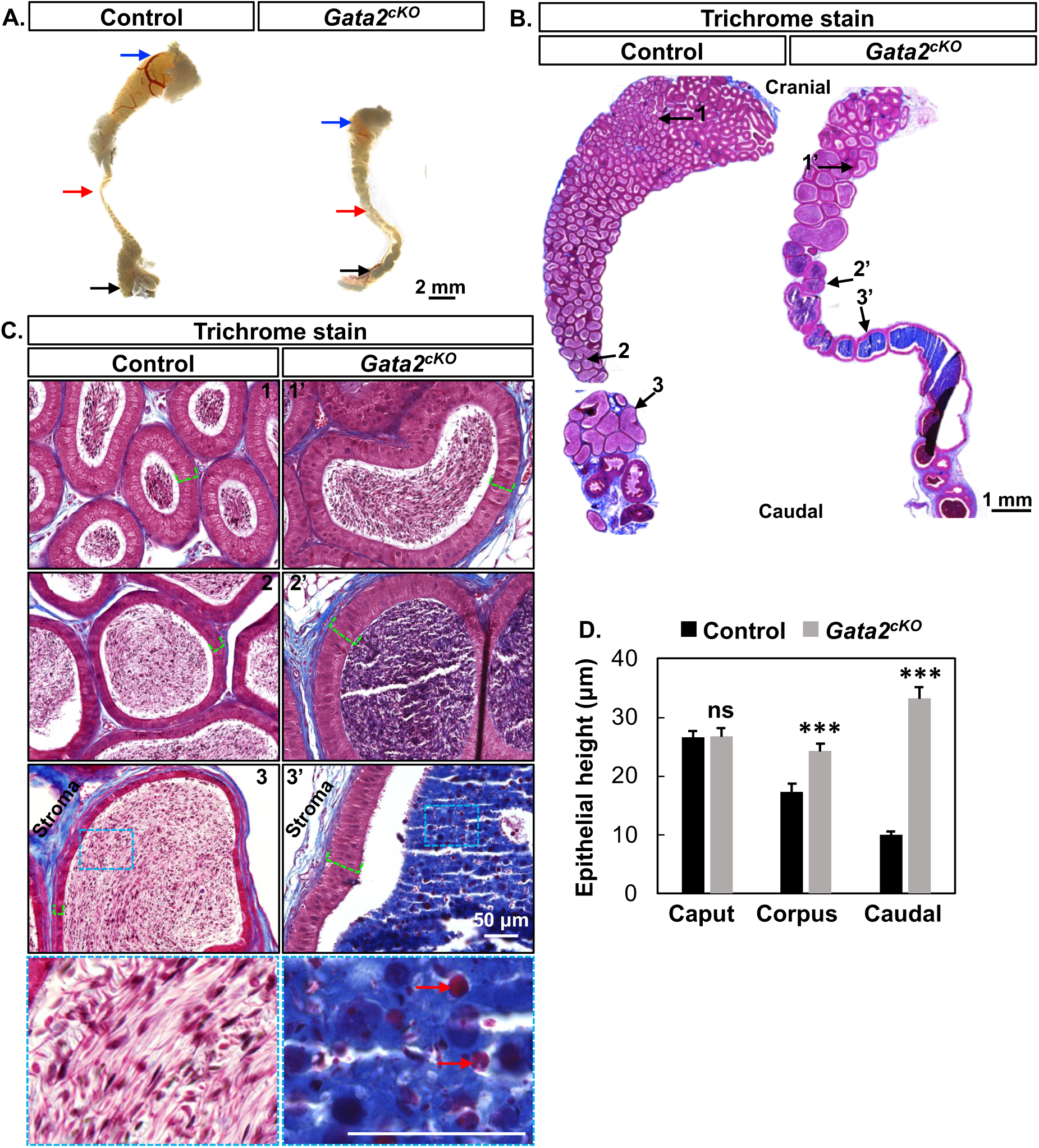
Adverse consequences on differentiation and function of the adult epididymis in *Gata2^cKO^* males. **(A)** Bright-field images of the control and *Gata2^cKO^* epididymides without testis at 8-week-old age with n=3 in each group. Blue, red and black arrows point to cranial, corpus, caudal regions, respectively. **(B)** Gomori trichrome staining on longitudinal sections of the control and *Gata2^cKO^* epididymides at 8-weeks old with n=6 in each group. 1-3 and 1’-3’ arrows indicated selected areas for higher magnification imaging in C. **(C)** Higher magnification images of cranial, corpus and caudal regions indicated by 1-3 arrows in the control and 1’-3’ in *Gata2^cKO^* epididymides in B with n=6 in each group. Green dashed lines indicate epithelial heights. Inserts in 3 and 3’: areas enlarged for visualization of lumen contents in the caudal epididymis. Red arrows: red blood cells. **(D)** Quantification of epithelial height in the cranial, corpus, and caudal regions between the control and *Gata2^cKO^*epididymides with n=3 males in each group. The average measurement of ten epithelia per region in each mouse were used for the statistical analysis. Results were shown as mean ± SEM and analyzed by 2-tailed *t* test. ns = not significant; *** *P* < 0.001.

To determine any histological differences between control and *Gata2^cKO^*epididymides, we used their longitudinal sections to perform Gomori trichrome staining, which allows for greater distinction between nuclear (black), cytoplasmic/muscle fibers (red/pink), and collogens/connective tissues (blue) (24) **(Fig. 4B)**. We observed multiple striking differences. First, the control epididymides showed extensive coiling throughout and displayed region-specific shapes: a club-shaped caput region, a narrower corpus region, and a wider caudal region. On the contrary, the epididymal coiling in the *Gata2^cKO^*male was reduced in the caput and completely absent in the corpus and caudal regions; and the shape of the caput, corpus and caudal regions remained relatively similar **(Fig. 4B)**. Another difference lay in the epithelial morphology and height. In control epididymides, epithelial cells were pseudostratified columnar in the cranial region and gradually transition to cuboidal epithelia in the caudal region. In control epididymides, the epithelial height gradually decreased from the cranial to the caudal regions (**Fig. 4C & 4D).** However, in the *Gata2^cKO^* epididymis, epithelial cells remained pseudostratified columnar cells in all the caput, corpus and caudal regions **(Fig. 4C)**; and the epithelial height was significantly increased in the corpus and caudal regions compared to that in the control epididymis **(Fig. 4D)**. Lastly, although sperm was detected in the lumen of the caput, corpus and caudal regions of both control and *Gata2^cKO^* epididymides, significant differences were observed in luminal contents between them. Strong blue staining indicative of collagens normally found in the stroma was detected in the lumen of corpus and caudal epididymides in *Gata2^cKO^* males **(Fig. 4B & 4C)**. Red blood cells, which were scarce in the lumen of control epididymides, became abundant in the lumen of *Gata2^cKO^* epididymides. These observations suggested that the lumen environment was altered in the *Gata2^cKO^* corpus/caudal epididymides and became unfavorable for sperm maturation and storage. In conclusion, our results demonstrated that the deletion of mesenchymal *Gata2* has long-term adverse effects on epididymal development and function.

## Discussion

### Mesenchymal GATA2 governs epididymal morphogenesis

The embryonic lethality of the global *Gata2* knockout at E10.5 due to defective hematopoiesis limited our investigations into GATA2’s roles in reproductive organ development at later stages (15). When *Gata2* expression was restored in hematopoietic development but remained absent in the whole urogenital system, the compound *Gata2* mutant embryos displayed fused vas deferens and seminal vesicles (25). However, the mesenchyme-specific function of *Gata2* during epididymal development remains unknown. In this study, we used a mesenchyme-specific conditional knockout mouse model to reveal the previously unrecognized role of mesenchymal *Gata2* promoting epididymal coiling. Mesenchymal GATA2 has been reported to play important roles in the morphogenesis of several other tubular organs (11–13, 26). For example, transgenic complementation of mesenchymal *Gata2* expression rescued improper displacement of ureter budding in *Gata2* hypomorphic mutant mice (26). In another study, conditional deletion of *Gata2* in the mesenchyme using *Tbx18-Cre* resulted in hydroureter formation at birth (12). In both models, the congenital anomalies of the kidney and the urinary tract led to neonatal and early postnatal deaths (27). The *Osr2-Cre* knock-in line used in our study is not active in the ureteric bud structure and does not exhibit ectopic Cre activity (27, 28). As a result, *Gata2^cKO^* mice did not develop any congenital anomalies in the kidney and the urinary tract and were able to survive to adulthood in our study.

Our discovery of the critical role of mesenchymal *Gata2* echoes classical tissue recombination experiments, which demonstrate the critical role of the mesenchyme in instructing WD morphogenesis. When the upper WD epithelium (future epididymis) was combined with the mesenchyme of the lower WD (future seminal vesicle), the upper WD epithelium was reprogrammed towards seminal vesicle-like morphogenesis and differentiation (8). It is well-established that *Hox* genes, mesenchymal transcription factors expressed along the craniocaudal axis, determine positional identity and morphogenesis of WD epithelium (29). *Hoxa10* or *Hoxa11* deletion resulted in the cranial transformation of the cauda epididymis and the proximal vas deferens, which became highly coiled (30–32). In contrast, the corpus and caudal epididymides in our *Gata2^cKO^* model underwent the caudal transformation into a straight and vas deferens- like tube. Our results indicate that in normal conditions, the function of mesenchymal GATA2 is to protect the epididymis from shifting toward the vas deferens. This discovery prompts us to propose a new concept in epididymal development. To establish the fate and morphogenesis of the middle WD, its transformation into the cranial and caudal regions need to be blocked by HOXA10/11 and GATA2, respectively. Further studies will be needed to understand how HOXA10/11 and GATA2 antagonize each other’s action in determining epididymal fate and differentiation.

### Components in androgen signaling are unaffected in *Gata2^cKO^*male embryos

Epididymal development has been predominantly linked with AR signaling (1, 18). Male mice embryos with a deficiency in *Cyp17a1,* a key steroidogenic enzyme in androgen synthesis, lose their epididymides (33). In utero exposure to AR antagonists, such as flutamide, significantly decreased epididymal coiling (17). Therefore, we initially speculated that defective epididymal coiling in the *Gata2^cKO^* male embryos resulted from deficient androgen signaling. However, *Gata2* is not expressed in the fetal testis (34), and we did not observe any changes in the expression of the key steroidogenic enzymes in the testis. In prostate cancer cell lines, *Ar* expression is promoted by GATA2 while GATA2 expression is repressed by androgen-AR action, forming a negative feedback regulatory loop (35). However, in our study, mesonephric AR expression was not altered in the absence of mesenchymal *Gata2*; and GATA2/*Gata2* expression was not changed in the absence of mesenchymal *Ar*. This discrepancy may be due to fundamental differences in gene regulatory networks between normal development and pathological cancer conditions. The defective epididymal coiling, despite normal androgen production and AR expression, suggests that androgen signaling alone is not sufficient to facilitate epididymal coiling and that local signaling factors are also indispensable for epididymal development.

It is noteworthy that GATA2 has been identified as a “pioneer factor” for AR in the prostate cancer epithelial cells, facilitating chromatin accessibility for AR binding and transcriptional regulation (36, 37). GATA2 occupies more than half of AR binding sites, and *Gata2* deletion dramatically impacts AR genomic distribution (36, 37).Therefore, one immediate follow-up question is whether mesenchymal GATA2 serves as a pioneer factor for mesenchymal AR in promoting WD development. In this study, we did not provide mesenchyme-specific genome-wide bindings of GATA2 and AR. However, multiple lines of indirect evidence suggest that mesenchymal GATA2 may not function as a pioneer factor for mesenchymal AR in the context of WD development. First, deletion of *Gata2* and *Ar* in the mesenchyme using the same Cre resulted in different phenotypes (defective coiling vs degeneration) (10), indicating distinct developmental gene programs regulated by these two transcription factors. Furthermore, our chromatin accessibility data on WD mesenchyme during sexual differentiation revealed enrichment of AR motif but not GATA motif in male-specific open chromatin regions (14). Lastly, AR ChIP-seq on adult epididymides did not identify the enrichment of GATA motifs in AR binding sites (38). Therefore, we suspect that the pioneer factor nature of GATA2 for AR observed in prostate cancer cells does not apply to the mesonephric mesenchyme during WD development.

### *Inhba* is a potential downstream effector of mesenchymal GATA2

An important finding of our study is that mesenchymal GATA2 is a key transcriptional regulator of *Inhba* expression. *Inhba* is the only known mesenchyme-derived secreted factor that plays a crucial role in epididymal coiling (21). *Inhba* expression is first present in the cranial region of the mesonephroi in both male and female E12.5 embryos, indicating that its initial mesenchyme-specific expression is not male-specific and does not require androgen signaling. From E13.5 onward, the testis is required for *Inhba* expression, which can be partially induced by testosterone. This observation suggests that other factor(s) from the testis may exist in maintaining *Inhba* expression after E13.5. Since *Gata2* is not expressed in the fetal testis, and we did not observe any defects in the testicular development, we hypothesize that testicular factors did not contribute to the decreased expression of *Inhba* in *Gata2^cKO^* males. Instead, the local mesenchymal *Gata2* may directly regulate *Inhba*, which has been shown to be a GATA2 target gene in mouse embryonic stem cell lines based on ENCODE Transcription Factor Targets Dataset (39). Further studies are needed to determine whether *Inhba* is indeed a direct GATA2 target gene in the WD mesenchyme.

The major product of *Inhba* in the mesonephros has been suggested to be activin A (homodimer of *Inhba*) that promotes WD epithelial proliferation (21). Activin A belongs to the TGFβ/BMP family of ligands and binds to its cognate type I receptor (*Acvr1*, *Acvr1b*, or *Acvr1c*) and II receptors (*Acvr2a* or *Acvr2b*), which can lead to the phosphorylation of type I receptor and ultimately activate SMAD-dependent and independent signaling in the context of target cells (22, 40). WD epithelium expresses Activin A receptors (type I *Acvr1b* and type II Acvr2a and *Acvr2b*) (41, 42) and the intracellular effector p-SMAD2/3, suggesting active activin A-SMAD signaling (21).

Since the common mediator, SMAD4, is a central component of the intracellular SMAD complex, we originally expected that deletion of *Smad4* in WD epithelium could result in defective epididymal coiling, phenocopying the *Inhba* knockout phenotype. However, the epididymis underwent normal coiling upon the deletion of epithelial *Smad4* using *Hoxb7-Cre*. *Hoxb7-Cre* is active as early as E10.5 in the WD epithelium, but its Cre recombination only occurs in approximately 60-80% WD epithelium (43). Consistent with the report, we also observed SMAD4 expression in a portion of epithelial cells of *Smad4^cKO^* WDs. Therefore, the lack of any morphological phenotype might be due to incomplete Cre recombination. Because activin A activates SMAD-dependent and independent signaling in the context of target cells (22, 40), another explanation is that activin A may act through SMAD-independent intracellular pathways in promoting epididymal coiling.

### Epididymal defects in *Gata2^cKO^* males persist into adulthood

A significant strength of our mouse model is that *Gata2^cKO^*males can survive to adulthood without any gross defects, allowing us to investigate the long-term impacts of deleting mesenchymal *Gata2* on the adult epididymis. Because the epididymis continues to undergo extensive coiling after birth (7), it is possible that the uncoiled *Gata2^cKO^* epididymides at birth might catch up on the coiling during postnatal development. However, they remained straight and were unable to coil, highlighting the crucial role of fetal development in establishing proper epididymal morphology in adulthood. These results align with the notion proposed by Welsh et al. that morphogenesis and differentiation of male reproductive tract organs must take place within a specific fetal programming window (5, 6). Decreased epithelial cell height in the caput epididymis has been shown to indicate androgen deprivation (44). However, we observed normal epithelial height in the caput region of *Gata2^cKO^* adult epididymides.

Therefore, the persistent defects are likely not due to a lack of sufficient androgens. Deletion of mesenchymal *Gata2* might permanently alter the identity of organ stroma and stroma-derived instructive signal, as evidenced by the reprogramming of the cuboidal epithelium into pseudo-stratified columnar epithelium with increased heights in the caudal region. It remains unclear whether the defects observed in adulthood are the direct outcome of *Gata2* deletion and/or the indirect consequence of the lack of epididymal coiling. To determine the direct impact of deleting mesenchymal *Gata2* on postnatal epididymal function, we propose using tamoxifen inducible Cre models (such as *Wt1-Cr*e*ER* (45)) to delete *Gata2* at the adult stage, which could be an interesting future direction.

The abundant presence of immune cells in the lumen of the cauda epididymis in *Gata2^cKO^* adult males might suggest the disruption of the blood-epididymal barrier, which regulates the content of the epididymal lumen by limiting molecule and cell movement between the blood and lumen (46, 47). The loss of the barrier might expose sperm to the male immune system and trigger local immune responses. The most significant influx of immune cells and luminal content changes occurred in the caudal region, possibly due to the more pro-inflammatory environment in the cauda compared to the caput and corpus regions (24).

### Conclusion

Our study demonstrates the androgen-independent role of mesenchymal GATA2 in promoting epididymal development possibly by inducing mesenchyme-derived factor *Inhba*. The epididymal defects caused by the fetal loss of mesenchymal *Gata2* persist into adulthood, supporting the hypothesis of fetal origins of adult reproductive disorders in a larger context (48). As far as we know, our study was the first report of a critical mesenchymal transcription factor other than AR in promoting epididymal coiling. In humans, GATA2 is also expressed in the WD mesenchyme (49, 50), indicating the relevance of our work to human male reproductive tract development. Similar congenital defects of the epididymis, such as hypoplasia and partial agenesis, have been reported in humans (51, 52). Therefore, our work will not only significantly advance our understanding of normal WD development, but also potentially help the development of improved strategies for prevention, diagnosis, and treatment of abnormal male reproductive tract differentiation.

## Materials and Methods

### Mice

*Gata2-flox* mice were obtained from the University of Missouri Mutant Mouse Resource & Research Centers (MMRRC, 030290) on C57BL/6J background (53). *Osr2- Cre* (knock-in Cre) strain was obtained from Dr. Rulang Jiang at Cincinnati Children’s Hospital Medical Center on mixed genetic backgrounds (129/Sv X C57BL/6J) (28) and established in the University of Wisconsin-Madison (10). *Ar-flox* (018450), *Smad4-flox* on mixed genetic backgrounds (129S6/SvEvTac X C57BL/6) (017462), *Hoxb7-Cre* (004692) on C57BL/6 background were purchased from Jackson Laboratory. Timed mating was set up by housing two or three females (*flox/flox*) with a sexually mature male (*Cre+;flox/+*). Vaginal plugs were checked the next morning, and the day of the vaginal plug detection was designated as E0.5. *Osr2^Cre+^:Gata2^flox/flox^*was designated as *Gata2^cKO^* and genotypes *Osr2^Cre+^:Gata2^flox/+^*, *Osr2^Cre-^:Gata2^flox/flox^,* and *Osr2^Cre-^:Gata2^flox/+^*were designated as the control. *Osr2^Cre+^:Ar^flox^/Y* and *Osr2^Cre-^:Ar^flox^/Y* were designated as *Ar^cKO^*and as control, respectively. *Hoxb7^Cre+^:Smad4^flox/flox^* was designated as *Smad4^cKO^*while *Hoxb7^Cre+^:Smad4^flox/+^* , *Hoxb7^Cre-^:Smad4^flox/flox^* were designated as control. All mouse procedures were approved by the University of Wisconsin-Madison (UW-Madison) Animal Care and Use Committees and followed UW-Madison approved animal study proposals and public laws. The method of euthanasia used was CO2 inhalation followed by cervical dislocation for mice and decapitation for embryos, which is consistent with the American Veterinary Medical Association (AVMA) Guidelines for the Euthanasia of Animals.

### Genotyping

PCR was performed using primers for *Osr2-Cre* allele (Forward 5’- GTCCAATTTACTGACCGTACACC-3’, Reverse 5’-GTTATTCGGATCATCAGCTACACC-3’), *Gata2*-*flox* allele (Forward 5’- TGGTGCTTTTCCCTTCTGTGTCGA-3’, Reverse 5’-CCTTCCCTGTCCCCAACTTCTCCT-3’), *Ar-flox* allele (forward 5’- AAAATGCCTCCTTTTGAC CA-3’, reverse 5’-AAGATGACAGTCCCCACGAG-3’)*, Smad4-flox* allele (Forward 5’-TAAGAGCCACAGGGTCAAGC-3’, Reverse 5’- TTCCAGGAAAAACAGGGCTA-3’), and *Hoxb7-Cre* (Transgene 5’- GGTCACGTGGTCAGAAGAGG-3’, Transgene 5’-CTCATCACTCGTTGCATCGA-3’, Internal Positive Control Forward 5’-CAA ATGTTGCTTGTCTGGTG-3’, and Internal Positive Control Reverse 5’- GTCAGTCGAGTGCACAGTTT-3’). Thermal cycle used Platinum II Taq Hot-Start DNA Polymerase (Invitrogen, 14966001). The conditions for the thermal cycle were 94°C for 2 min, 34 cycles of [94°C for 15s, 60°C for 5s, and 68°C for 15s] followed by 68°C for 5 min.

### Tissue Processing, Embedding and Sectioning

We followed the procedure as previously described (54). Embedded tissues were fixed in 10% neutral buffered formalin (Leica, 3800598). Tissues underwent three 10-minute 1xPBS (phosphate-buffered saline) washes and followed a series of dehydration using ethanol (70%, 80%, and 95% for 30 min each; 100% ethanol I and II for 50 min each). After ethanol, tissues were then exposed to a 1:1 ethanol to xylene mixture followed by two xylene steps ranging from 5-10 minutes depending on sample size. The tissues were transferred to a 1:1 xylene to paraffin mixture for 10 minutes and followed three paraffin infiltrations for 1-2 hours each. All these steps were extended by 30 mins in processing adult tissues. For cross-sections, tissues from the cranial to caudal mesonephros/epididymis were sectioned at 5 μm using a microtome (Tanner Scientific, TN6000) with >50 μm apart between sections; for longitudinal sections, adult epididymides were sectioned at 5 μm with >30 μm apart.

### Histology

H&E (hematoxylin & eosin) staining was performed as previously described (54). Gomori Trichrome Staining Procedure No. HT10 was performed by the University of Wisconsin-Madison’s Histology Resource Core in the Department of Surgery using Trichrome Stain AB Solution (Millipore Sigma). After staining, all sections were imaged using a Keyence microscope.

### Quantifications of epithelial height

ImageJ was used to quantify the height of epithelial cells in 40x images with the width of 960 pixels, and the height of 720 pixels from the Keyence microscope. The software was first calibrated using 50 μm scale bar so 257 pixels were calibrated to equal 50 μm. The control and *Gata2^cKO^* groups had three mice each. In each mouse, ten epithelial heights from random tubules in the caput, corpus and caudal regions were measured. Results from these ten measurements were then averaged for the caput, corpus and caudal region for each mouse as a datapoint for statistical analysis.

### Immunofluorescent staining

We followed our established protocol as previously described (10). Briefly, all sections were subjected to antigen retrieval using antigen unmasking solution (VECTOR, catalog No. H-3300). After being heated during antigen retrieval, sections were cooled and transferred to Sequenza Immunostaining Center System (Thermo Fisher Scientific, catalog No. 73310017; cover plates: catalog No. 72110017). Sections were washed with 1×PBS and PBST (1× PBS with 0.1% Triton X- 100) then incubated in blocking buffer (5% normal donkey serum in 1× PBST) for 1 hour at room temperature followed by primary antibody incubation in blocking buffer at 4°C overnight. The following primary antibodies were used: Rabbit Anti-GATA2 (Millipore Sigma, HPA005633, 1:100); Rabbit Anti-ITGA4 (D2E1) XP (Cell signaling, 8440, 1:100); Rabbit Anti-KI67 (Abcam, ab15580, 1:200). After washing three times with PBST, sections were incubated with donkey anti-rabbit Alexa Fluor 488 conjugate (Invitrogen, A-21206,1:200) in blocking buffer for one hour at room temperature. Finally, sections were washed with PBST and counterstained with DAPI (Thermo Fisher Scientific, 62248, 1:1000) and cover-slipped with EverBrite mounting medium (Biotium, 23003) for imaging under a Leica TCS SP8 confocal microscope.

### Immunohistochemistry

We followed our previously established protocol (10). Paraffin sections were subjected to deparaffinization, rehydration and antigen retrieval (VECTOR, catalog No. H-3300). Sections were treated with 3% H2O2 to eliminate endogenous peroxidase activities, washed with PBST, and then incubated in the blocking buffer for one hour followed by incubation with primary antibody in the blocking buffer at 4 °C overnight. The primary antibodies included Rabbit Anti-Androgen Receptor (Abcam, 133273, 1:200) and Rabbit Anti-SMAD4 (Abcam, ab40759, 1:200). The next day, sections were washed with PBST three times and then incubated with biotinylated Donkey Anti-Rabbit IgG H&L (Abcam, ab97062, 1:200) at room temperature for 30 min. Sections were then incubated with ABComplex/HRP (VECTOR, PK-6100) and DAB substrate (VECTOR, SK-4100), counterstained with hematoxylin. All sections were then imaged using a Keyence microscope.

### RT-qPCR

E16.5 mesonephroi and testes were separated and snap-frozen on dry ice. RNA extraction and cDNA synthesis were performed using PicoPure RNA Isolation kit (Life technologies, KIT0204) and the Superscript cDNA synthesis kit (Qiagen, 330411), respectively, according to manufacturer’s protocols. For cDNA synthesis, 200ng of RNA in each sample was used. SYBR-green master mix was used (Thermo Fisher Scientific, 208052) with the following primers: *Hsd3b1*, 6680289a1 (Harvard PrimerBank ID); *Cyp17a1*, 6681097a1; *Gapdh*, 6679937a1; *Itga4*, 7110657a1; *Ar*, 7304901a1; and *Gata2* flanking exon 5 and 6 (Forward 5’- GCACCTGTTGTGCAAATTGT-3’, Reverse 5’- AGGGCGGTGACTTCTCTTG-3’) (55). All samples were run in duplicates and normalized to the housekeep gene *Gapdh*. The relative expression in the knockout group was reported as a ratio of the expression relative to those of control mice.

### RNAscope and quantification of *Inhba* staining

RNAscope was performed according to the manufacturer’s protocol (14). After deparaffinization and rehydration, paraffin sections were treated with hydrogen peroxide, antigen retrieval buffer and proteinase. Sections were then incubated with *Inhba* probe (Advanced Cell Diagnostics, 455871) and followed a series of rinsing and signaling detection at designated temperatures. All sections were then imaged using a Keyence microscope. ImageJ was used to quantify the expression of *Inhba* in the mesenchyme. There were four control and four knockout mice each with four to eight cross-sections from the cranial to the caudal region. The sections were >50 μm apart between. To quantify the percentage of *Inhba+* cells, the layer of mesenchymal cells most adjacent to the epithelium in each section was used to calculate the fraction of *Inhba+* cells over the total number of adjacent mesenchymal cells. To quantify the average expression intensity in those *Inhba+* mesenchymal cells, we used Advanced Cell Diagnostics (ACD) semi-quantitative scoring system: 1 = 1-3 dots/cell, 2 = 4-9 dots/cell with very few clusters, 3 = 10-15 dots/cell with <10% of dots in clusters, and 4 = >15 dots/cell and >10% of dots in clusters. Results of four to eight sections from each mouse were averaged as a data point for statistical analysis.

### Quantifications of KI67+ cells

Mesonephroi from six embryos in each group were sectioned from the cranial to the caudal regions. For each mesonephros, four to eight sections with >50 μm apart between sections were used for KI67 staining and subsequent quantification. In each section, the percentage of KI67+ cells was calculated as the fraction of KI67+ epithelial cells over the total number of epithelial cells (identified by DAPI nuclei staining). The results in all the sections from the same mesonephros were averaged as a datapoint for statistical analysis.

### Organ culture

E16.5 mesonephroi with testes attached were cultured on MilliCELL-CM culture insert 0.4 µm filters (MilliporeSigma, PICM03050) in DMEM/F12 (Gibco, 21041025) supplemented with 100 U/mL penicillin–streptomycin (Gibco, 15070063) at 37°C in a 5% CO2 incubator (20). Organ cultures were supplemented with 10nM dihydrotestosterone (Selleck Chemicals, S4757) or no treatment. Culture media was changed every other day. Bright field images were taken daily with a Leica S9 scope.

### Statistical analyses

A minimum of three biological replicates were used in all experiments. The sample sizes in each data set were specified in figure legends. The significant differences between control and knockout groups were evaluated by two-tailed unpaired Student’s t-test. Results were shown as mean ± SEM. The significance level was set at *P* < 0.05.

## Supporting information

Supplementary figures with figure legends

## Acknowledgements.

The authors thank the University of Wisconsin-Madison, Department of Surgery Histology Resource Core (HRC), Dr. Angela Gibson, MD, PhD, FACS, PI of the HRC, along with certified histotechnologist Sierra Raglin, HTL (ASCP) CMP and Rebecca Lyons-Oelze, HTL (ASCP) for performing trichrome staining. We would also like to thank Dr. Rulang Jiang from Cincinnati Children’s Hospital Medical Center, Division of Developmental Biology for providing *Osr2-Cre* mice, Dr. McDowell, DOVS, from the University of Wisconsin-Madison for providing an aliquot of ITGA4 antibody, and Mutant Mouse Regional Resource Center U430D010918 for providing *Gata2-flox* mice.

## Data availability

All data are available in the main text or the supplementary materials upon reasonable request.

## Author contributions

Allyssa Fogarty: Writing original draft and review & editing of final draft; Conceptualization; Formal analysis; Investigation; Project administration; Visualization. Shuai Jia: Investigation; Visualisation; Validation; review & editing of final draft. Jillian Wilbourne: Investigation; review & editing of final draft. Claire DuPuis: Investigation; review & editing of final draft. Fei Zhao: Conceptualization; Methodology; Writing original draft and review & editing of final draft; Supervision; Funding acquisition.

