## Supplementary figures with figure legends for "Crucial roles of mesenchymal *Gata2* in murine epididymal development"

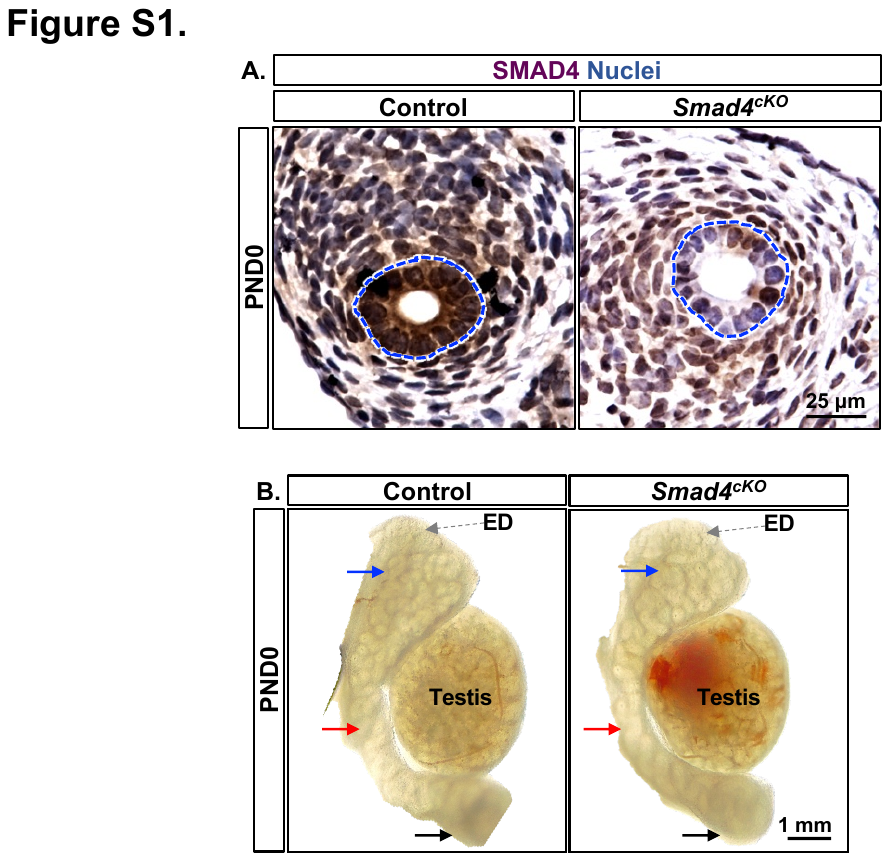
**Supplementary Figure 1.** Deletion of epithelial *Smad4* did not result in observable phenotypes. **(A)** Immunohistochemistry staining of SMAD4 on the cross-sections of control and *Smad4^cKO^* epididymides with n=4 in each group, 2 at PND0 and 2 at PND21. Blue dashed lines: WDs. **(B)** Bright-field images of the control and *Smad4^cKO^* epididymides with n=4 in each group, 2 at PND0 2 at PND21. Blue, red and black arrows indicate cranial, corpus, and caudal regions, respectively. The grey arrows: ED.

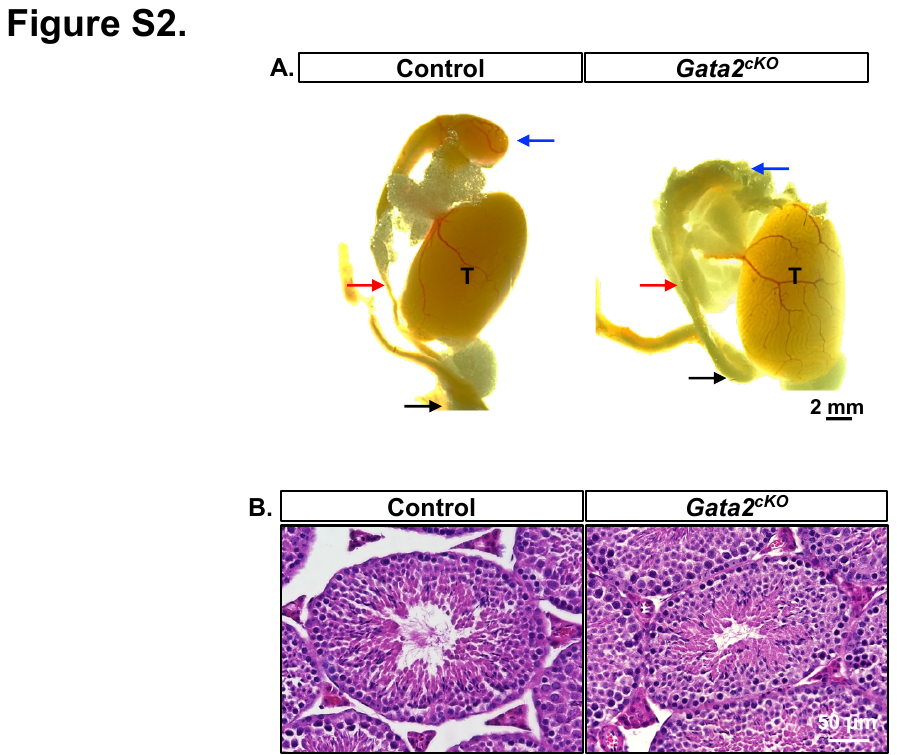

**Supplementary Figure 2.** Morphology of the adult epididymides and testes in *Gata2^cKO^* males. **(A)** Bright-field images of the control and *Gata2^cKO^* epididymides with testis at 8-weeks with n=3 males in each group. **(B)** Hematoxylin and Eosin staining of the control and *Gata2^cKO^* testes at 8-weeks with n=3 males in each group.
